# Hippocampal hub failure is linked to long-term memory impairment in anti-NMDA-receptor encephalitis - Insights from structural connectome graph theoretical network analysis

**DOI:** 10.1101/2023.08.18.553940

**Authors:** André Hechler, Joseph Kuchling, Leonie Müller-Jensen, Johanna Klag, Friedemann Paul, Harald Prüss, Carsten Finke

**Author notes:** **Correspondence:** Carsten Finke, Department of Neurology, Charité - Universitätsmedizin Berlin, Charitéplatz 1, 10117 Berlin, Germany. Equally contributing first authors (André Hechler, Joseph Kuchling).

## Abstract

**Introduction:** Anti-N-methyl-D-aspartate receptor (NMDAR) encephalitis is characterized by distinct structural and functional brain alterations, predominantly affecting the medial temporal lobes and the hippocampus. Structural connectome analysis with graph-based investigations of network properties allows for an in-depth characterization of global and local network changes and their relationship with clinical deficits in NMDAR encephalitis.

**Objective:** To investigate changes in structural connectivity and network efficiency in NMDAR encephalitis by use of probabilistic whole-brain tractography and graph theoretical analysis of structural brain networks.

**Methods:** Structural networks from sixty-one NMDAR encephalitis patients in the post-acute stage (median time from acute hospital discharge: 18 months) and sixty-one age- and sex-matched healthy controls (HC) were analyzed using diffusion-weighted imaging (DWI)-based probabilistic anatomically-constrained tractography and spherical deconvolution-informed filtering of tractograms. We calculated global, modular, and nodal graph measures indicative of structural connectivity and network reorganization with special focus on default-mode network, medial temporal lobe, and hippocampus. Pathologically altered metrics were included in multiple regression analyses to investigate their potential association with clinical course, disease severity, and cognitive outcome.

**Results:** Patients with NMDAR encephalitis showed regular global graph metrics, but bilateral reductions of hippocampal node strength (left: *p*=0.049; right: *p*=0.013) and increased node strength of right precuneus (*p*=0.013) compared to HC. Betweenness centrality was decreased for left-sided entorhinal cortex (*p*=0.042) and left caudal middle frontal gyrus (p = 0.037). Correlation analyses showed a significant association between reduced left hippocampal node strength and verbal long-term memory impairment (*p*=0.021)

**Conclusion:** Focal network property changes of the medial temporal lobes indicate hippocampal hub failure that is associated with memory impairment in NMDAR encephalitis at the post-acute stage, while global structural network properties remain unaltered. Graph theory analysis provides new pathophysiological insight into structural network changes and their association with persistent cognitive deficits in NMDAR encephalitis.

## Introduction

Anti-N-methyl-D-aspartate receptor (NMDAR) encephalitis is an autoimmune encephalitis with a characteristisc neuropsychiatric syndrome that can include behavioral changes, movement disorders, hallucinations, seizures and cognitive deficits [1,2]. Despite the frequently severe clinical course [3], the individual level of disease burden in NMDAR encephalitis is not reflected by the degree of abnormalities in routine MRI, which is unremarkable in 50-77% of patients [4].

By contrast, functional MRI analyses showed impaired connectivity between the hippocampus and the default mode network (DMN) that correlated with memory impairment [5]. Investigations of brain-wide functional networks demonstrated both wide-spread changes in fronto-medial and fronto-parietal connections and focal disruptions within the medial temporal lobe (MTL) network, with the latter being closely associated with disease severity and memory performance [6]. Moreover, diffusion tensor imaging (DTI) investigations also revealed persistent structural damage. For example, widespread alterations of fractional anisotropy (FA) and mean diffusivity (MD) that correlated with disease severity, reflecting profound white matter damage in NMDAR encephalitis [5]. In addition, brain-wide alterations in superficial white-matter diffusivity were observed and were associated with disease severity and with persistent deficits of working memory, verbal memory, visuospatial memory and attention [7]. Interestingly, patient-derived NMDAR antibodies have recently been shown to alter NMDA receptor function in oligodendrocytes, suggesting a link between antibody-mediated dysfunction of NMDARs in oligodendrocytes and white matter alterations detected using MRI analyses [8]. However, despite these functional network changes and structural white matter alterations, detailed investigations of structural connectivity and network efficiency changes caused by NMDAR encephalitis and their potential association with clinical and cognitive deficits are still missing.

Diffusion-weighted imaging (DWI) based probabilistic tractography allows for a high resolution reconstruction of white matter tracts within the whole brain [9]. In recent years, advanced tractography algorithms have been developed based on methods such as constrained spherical deconvolution (CSD) [10,11] and *a priori* tract boundary knowledge implemented in anatomically-constrained tractography (ACT) [12]. Moreover, graph theory analysis has emerged as a key tool to derive topology motifs from structural brain connectivity datasets generated using probabilistic tractography [13]. These analyses have consistently identified central brain regions (“hubs”) that are critically important for efficient brain communication given their role for the integration of distributed neural activity [14]. However, their high level of centrality also renders hubs particularly susceptible to disconnection and dysfunction. Indeed, graph theoretical analyses have identified dysfunction of a set of network parameters to be closely related to clinical and cognitive symptoms in multiple sclerosis (MS) [15,16], neuromyelitis optica spectrum disorders [17], schizophrenia [18], and Alzheimer’s disease [19,20].

Here, we aimed to generate structural networks using an analysis pipeline of CSD-based probabilistic tractography within the ACT framework [10–12] to evaluate global, modular and nodal characteristics of structural networks from NMDAR encephalitis patients. We then correlated pathologically altered network metrics with clinical and cognitive measures to elucidate potential associations between structural network changes and clinical disability.

## Methods

### Participants

Sixty-one patients with NMDAR encephalitis in the post-acute stage (52 [86.9%] female patients; median age = 25 years [range: 15 – 49 years]; median time from acute hospital discharge: 18 months; for further data see Table 1) were recruited from the Department of Neurology at Charité – Universitätsmedizin Berlin. Investigations comprised clinical evaluation, comprehensive neuropsychological assessment, and magnetic resonance imaging (MRI) data acquisition. Characteristic clinical presentation and detection of IgG NMDA receptor antibodies in the cerebrospinal fluid served as the basis for diagnosis according to current guidelines [2]. We enrolled 61 age- and sex-matched healthy participants without neurological or psychiatric diseases from our ongoing imaging database (EA4/011/19 and EA 1/163/12) to serve as healthy controls (HC). All participants gave written informed consent for all investigations and scientific publication of data prior to their inclusion in the study. The study was approved by the Charité ethics committee (EA4/011/19) and was performed in accordance with The Code of Ethics of the World Medical Association (1964 Declaration of Helsinki) in its currently applicable version.

**Table 1.** Clinical cohort description

|  | **NMDAR encephalitis** | **Healthy Controls** |
| --- | --- | --- |
| **Sex, n (% female)** | 53 female / 8 male<br>[86.9%] | 52 female / 9 male<br>[85.2%] |
| **Age, yr, median ± SD (range)** | 25.5 ± 8.9 (15-49) | 26.5 ± 8.7 (16-51) |
| **Onset-treatment interval [d; median (IQR)]** | 19 (100) | - |
| **Time since discharge from hospital [m; median (IQR)]** | 18.0 (24.0) | - |
| **Acute-stage mRS [median (range)]** | 4 (2-5) | - |
| **mRS at MRI [median (range)]** | 1 (0-3) | - |
| **Tumor [n (%)]** | 12 (19.7%) | - |

Abbreviations: IQR = interquartile range; SD = standard deviation; m = months; yr = years; d = days; mRS = modified Rankin scale.

### Neuropsychological assessment

All NMDAR encephalitis patients underwent comprehensive neuropsychological assessment for attention using the Test of Attentional Performance (TAP Version 2.3.1) [21], verbal memory using the Rey Auditory Verbal Learning Test (RAVLT) [22,23], spatial memory using the Rey-Osterrieth Complex Figure Test (ROCF) [24,25] and executive function using the Stroop Color and Word Test (SCWT) [26], as described in detail previously [27].

### MRI acquisition

All MRI data were acquired on the same 3T scanner (Tim Trio Siemens, Erlangen, Germany) using a single-shot echo planar imaging sequence for diffusion MRI acquisition (repetition time [TR] = 7500 ms, echo time [TE] = 86 ms; field of view [FOV] = 240 × 240 mm ; voxel size = 2.5×2.5×2.3 mm3, 61 slices, 64 non-colinear directions, b-value = 1000 s/mm) and a volumetric high-resolution T1 weighted magnetization prepared rapid acquisition gradient echo (MPRAGE) sequence (TR/TE/inversion time [TI] = 1900/2.55/900 ms, FOV = 240 × 240 mm2, matrix size = 240 × 240, 176 slices, slice thickness = 1 mm).

### DWI preprocessing and anatomically-constrained probabilistic tractography

DWI preprocessing and probabilistic tractography were performed using MRtrix3 [28], FMRIB Software Library’s (FSL) [29] and Advanced Normalization Tools (ANTs) [30], according to a previously described protocol [12,31] (Figure 1). The preprocessing included denoising, eddy-current correction, motion correction (using FSL topup) and bias-field correction (using ANTs). Structural T1 scans were parcellated into a total of 84 cortical and subcortical areas using the standard Freesurfer pipeline [32]. Tissue segmentation was performed using FSL FAST [33] as implemented in MRtrix3. Local fODF were obtained with probabilistic tractography using single-tissue CSD [10]. For improved streamline trajectories and rejection, we used ACT and limited seeding and termination to the interface of white matter and cortical or subcortical grey matter based on the segmented anatomical image [12].

**Figure 1.**
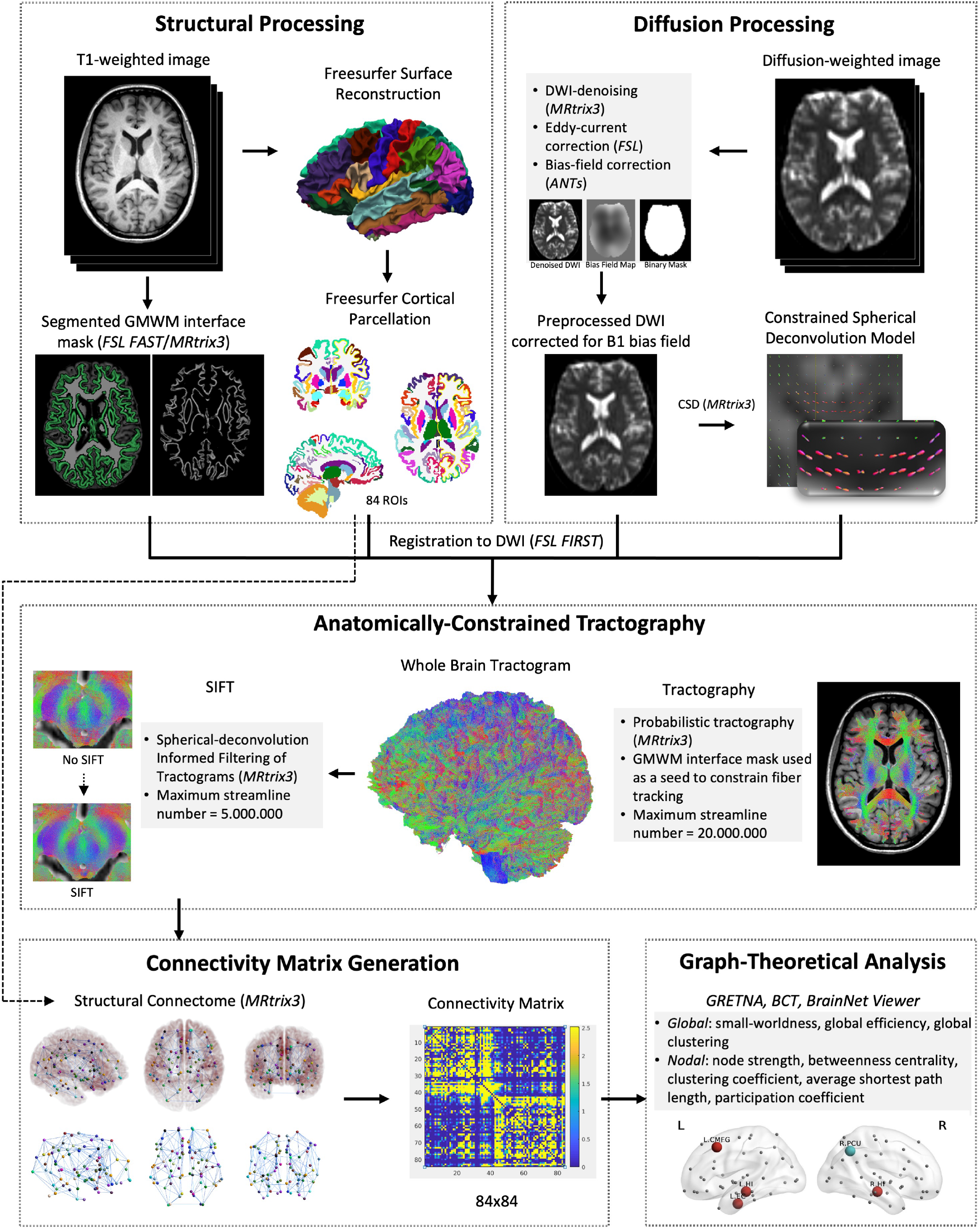
MR data and structural connectome processing pipeline. Structural T1 scans were parcellated into a total of 84 cortical and subcortical areas. Tissue segmentation was performed using FSL FAST [33] to generate a gray-matter white-matter (GMWM) interface mask as implemented in MRtrix3. Diffusion-weighted images (DWI) preprocessing included denoising, eddy-current correction, motion correction and bias-field correction. Local fiber orientation density functions (fODF) were obtained using single-tissue CSD [10]. For improved streamline trajectories and rejection, we used anatomically-constrained tractography (ACT) [12]. We then filtered the tractogram using spherical deconvolution-based filtering of tractograms (SIFT) [34]. Structural connectivity matrices were created based on the results of ACT and SIFT with columns and rows corresponding to the 84 anatomical regions (nodes) and cells corresponding to the number of streamlines (edges) between pairs of nodes. We used the Graph Theoretical Network Analysis Toolbox (GRETNA) [35] and the Brain Connectivity Toolbox (BCT) [36] to carry out graph analyses. Graphs were visualized using BrainNet Viewer [37]. Small-worldness, global efficiency and global clustering coefficient were calculated for whole networks and node strength, betweenness centrality (BC), clustering coefficient (CC), average shortest path length (APL) and participation coefficient (PC) for all nodes.

We then filtered the tractogram using SIFT [34], which has been shown to improve biological plausibility of the reconstructed tracts by discarding streamlines that do not correspond well to the underlying diffusion signal [38]. To balance the risk of an overabundance of false positive fibers (weak filtering) against the risk of artificially sparse tractograms (strong filtering) and to address limitations of computational demand (large amounts of streamline creation or strong filtering), an overall 20 million streamlines were created and subsequently filtered down to 5 million streamlines, gaining connectome accuracy comparable to previously published literature [12,34,38].

### Graph theory-based network analysis

Structural connectivity matrices were created based on the results of ACT and SIFT with columns and rows corresponding to the 84 anatomical regions (nodes) and cells corresponding to the number of streamlines (edges) between pairs of nodes. We used the Graph-Theoretical Network Analysis Toolbox (GRETNA) [35] and the Brain Connectivity Toolbox (BCT) [36] to carry out graph analyses (see Figure 1). Graphs were visualized using BrainNet Viewer [37]. Raw connectivity matrices contained a high number of edges with low probability, resulting in very dense networks that can distort classical graph measures [39]. Therefore, we integrated graph measures over a range of cutoffs (1% and 10% wiring cost in steps of 1% and thresholds between 10% and 90% in steps of 5%) as suggested previously [40]. A range of ten thresholds in steps of 5% connection density was chosen, with the upper bound defined as the most liberal threshold resulting in consistent small-worldness and the lower bound defined as the most conservative threshold that did not result in complete fragmentation of nodes. In our data, small-worldness (indicated by Sigma) showed marked variation over the complete range of thresholds (1% – 90% connection density) but low between-subject variability on individual levels (see Table 2). As networks lost consistent small-worldness upwards of 55%, this was defined as the upper bound of the threshold range. For networks below 10% connection density, fragmented nodes (node strength of 0) occurred with increasing probability. Consequently, we chose 10% as the lower bound. All tests on graph metrics were carried out using the area under the curve (AuC) over the described threshold range.

We subsequently tested differences in nodal graph metrics on Freesurfer based parcellation areas pertaining to the MTL (hippocampus, parahippocampal gyrus and entorhinal cortex) [41] and DMN (bilateral medial orbitofrontal gyrus, caudal medial frontal gyrus, caudal and rostral anterior cingulate, posterior cingulate, precuneus and inferior parietal gyrus) [42] since we expected marked network changes most likely to occur in these regions based on MR alterations observed in previous studies [5,6,43]. Small-worldness, global efficiency and global clustering coefficient were calculated for whole networks and node strength, betweenness centrality (BC), clustering coefficient (CC), average shortest path length (APL) and participation coefficient (PC) for all nodes. Small-worldness was derived from the sigma coefficient with values above 1 indicating small-world properties. Network hubs were defined using a hub score based on ranking all nodes on node strength, BC, APL and PC. Nodes falling into the top 25% in at least three of four measures were considered network hubs [14,39]. Nodes belonging to the MTL and DMN were tested for differences in node strength, BC, and CC. We additionally tested the connectivity within MTL and DMN by total node strength and mean APL.

### Statistical analysis

Subsequent statistical analyses were performed using R Studio (RStudio Team, 2015). Group differences in nodal and modular parameters between patients and HC were tested with non-parametric resampling (10,000 iterations) using the resample package [44]. Graph parameters showing significant differences were included in linear mixed-effects model analyses with age and years of education as covariates to investigate potential correlations with clinical (acute-stage mRS [45], onset-treatment-interval) and neuropsychological (RAVLT delayed recall, ROCF delayed recall,TAP go/no-go test) parameters. For all statistical analyses, a p-value of <0.05 was regarded as significant. Due to the exploratory nature of group comparisons and correlation analyses, we refrained from correction for multiple testing [46].

## Results

### Nodal graph metrics: MTL and DMN

Node strength in patients was significantly reduced in both left (p = 0.049) and right (p = 0.013) hippocampus relative to controls (Figure 2). A significant node strength increase was found within the right precuneus (p = 0.013). Differences in BC were found for the left caudal middle frontal gyrus (p = 0.037) and the left entorhinal cortex (p = 0.042), with lower values in NMDAR encephalitis patients (Figure 2). In addition, NMDAR encephalitis patients showed increased average path for left hippocampus (p = 0.017) and right parahippocampal gyrus (p = 0.026), while no differences were found with respect to the nodal CC. For a comprehensive overview of nodal graph metrics in DMN and MTL anatomical regions, see Supplementary Tables S 1 - 4.

**Figure 2.**
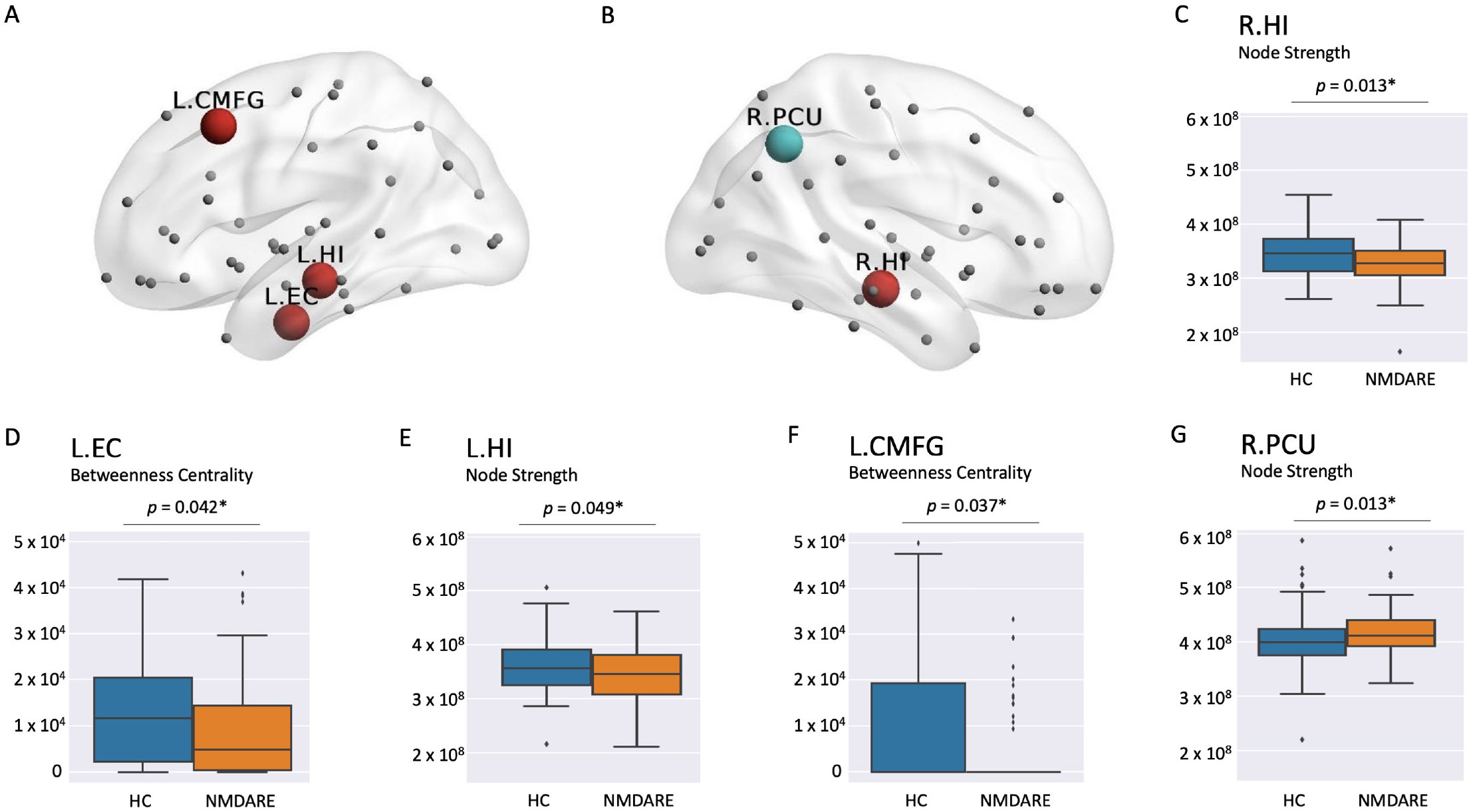
Group comparison of nodal graph metrics between NMDAR encephalitis and HC. **A** and **B** Lateral sagittal view with nodes exhibiting decreased (red) or increased (light blue) graph metrics in patients compared to HC. Red circles denote both decreases in node strength (L.HI, R.HI) or decreases in betweenness centrality (L.EC, L.CMFG). Light blue circles denote increases in node strength (R.PCU). **C – G** Comparative distribution of individual nodal graph metric values between HC (blue) and NMDAR encephalitis (orange) is displayed in the boxplots below for each region of interest. L.EC = left entorhinal cortex; L.HI = left hippocampus; L.CMFG = left caudal middle frontal gyrus; R.HI = right hippocampus; R.PCU = right precuneus

### Modular graph metrics: MTL and DMN

The MTL showed a non-significant trend towards lower values for patients in total node strength on the left (p = 0.09) and right (p = 0.08) side and mean APL on the left side (p = 0.05; Table 2). No marked differences were found for any graph metric in the DMN (see Supplementary Table S5).

### Global graph metrics

No significant differences in global efficiency and in global clustering coefficient calculation were found between NMDAR encephalitis patients and HC (see Supplementary Table S6).

### Correlation of graph metrics with clinical data

The following predictors were tested based on significant group differences: Node strength of the left and right hippocampus and the right precuneus as well as BC of the left entorhinal cortex and left caudal medial frontal cortex. Regarding clinical parameters, we found a significant association between left hippocampal node strength and verbal long-term memory (RAVLT delayed recall test; p = 0.021; Figure 3). No other associations or significant correlations were observed.

**Figure 3.**
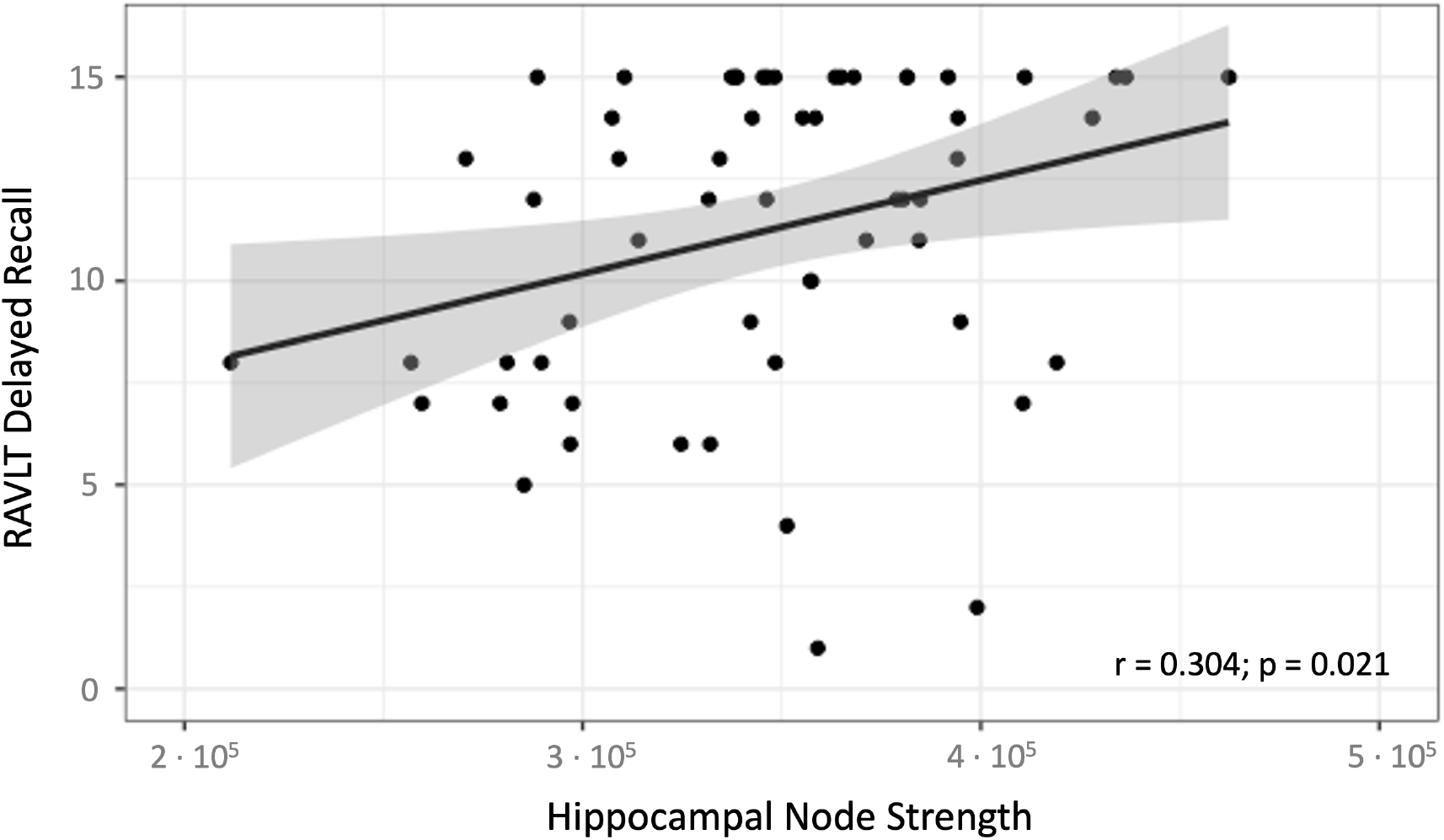
Association between left hippocampal node strength and verbal long-term memory. Scatter plot of multiple linear regression analysis with individual values of left hippocampal node strength and RAVLT delayed recall values that showed a strong positive correlation (regression line; r=0.304 [Pearson correlation coefficient]; p-value =0.021 [corrected for age and years of education]). RAVLT = Rey Auditory Verbal Learning Test

## Discussion

In this study, we investigated structural connectivity changes and multi-level network topology alterations in NMDAR encephalitis. To this end, we used an analysis framework with DWI based probabilistic and anatomically constrained tractography and integration of graph metrics over multiple thresholds. We observed reduced node strength in both hippocampi, but increased node strength in the right precuneus in NMDAR encephalitis patients compared to healthy controls, indicating structural network reorganization following hippocampal hub failure. On a modular subnetwork level, we detected decreased node strength in the left and right MTL and an increased path length for the left MTL in NMDAR encephalitis patients. Moreover, correlation analyses revealed an association between hippocampal node strength and verbal long-term memory in NMDAR encephalitis patients. By contrast, no significant differences in global network metrics were found between NMDAR encephalitis patients and HC. Overall, our study provides novel insights into structural connectivity disruptions at modular and nodal levels and the impact of this network disintegration pattern on individual disease burden in NMDAR encephalitis.

The MTL is the key brain structure related to episodic memory and damage to the MTL is associated with profound memory impairment [41,47,48]. Previous studies in NMDAR encephalitis patients reported on hippocampal atrophy and impaired microstructural integrity of the hippocampal formation that were correlated with memory deficits [5,49]. Our structural network findings with bilateral reductions in hippocampal node strength, reduced BC in the left entorhinal cortex, and increased APL in left hippocampus and right parahippocampal gyrus lend further support to the notion that hippocampal damage and associated structural network disruption play key roles in NMDAR encephalitis pathophysiology.

The connectivity architecture of the brain (the connectome) is characterized by a central core of highly interconnected hub regions that are critical for efficient communication [50]. These network hubs are brain regions with high node strength (i.e., high number of connections [edges] with other nodes in the network) and betweenness centrality (i.e., high number of shortest paths in a network that pass through this node) [36]. However, given their crucial importance, hubs are strategic vulnerability points and damage to hubs leads to extensive network disruption and prominent clinical symptoms [50]. The hippocampus is one of the key hubs in the connectome given its high node strength and betweenness centrality. Here, we observed a bilateral reduction of hippocampal node strength in patients with NMDAR encephalitis. These results thus show that the disease targets a central network region of the brain, leading to disrupted hub function of the hippocampus and impaired connectivity of this densely connected brain structure [14]. Interestingly, this is in line with observations that other neurological disorders, including Alzheimer’s disease, Parkinson’s disease and multiple sclerosis, are likewise associated with damage to highly connected hub nodes [20].

Our findings are also in line with previous reports on bilateral atrophy of the input and output regions of the hippocampal circuit alongside microstructural damage in both hippocampi in NMDAR encephalitis [43]. Recent resting-state functional MRI investigations provided evidence on wide-spread functional connectivity impairment within distributed large-scale functional networks, including sensorimotor, frontoparietal, lateral-temporal, and visual networks [5,6,51,52]. Here, we observed structural connectivity alterations affecting both hippocampi and areas outside MTL indicating that previously identified functional network changes might at least partially be based on these structural changes. These alterations are most likely caused by direct effects of anti-NMDAR antibodies on hippocampal neurons given their high density of NMDARs, eventually leading to substantial disruption of network topology with hippocampal hub failure [53].

Importantly, we observed a strong association between left hippocampal node strength and verbal long-term memory. These findings are in line with previous functional MRI investigations that reported on impaired functional connectivity between the hippocampus and the default mode network (DMN) that correlated with verbal memory impairment in NMDAR encephalitis [5] and reports on close associations between focal disruptions within the MTL functional network and verbal memory scores in NMDAR encephalitis [6]. Hence, structural and functional connectome based nodal hippocampal graph metrics might provide potential imaging markers of clinical relevance to assess cognitive status in NMDAR encephalitis in future studies. This is of particular importance since cognitive deficits are the main contributor to long-term morbidity in NMDAR encephalitis and may either improve or persist over time in the individual patient depending on yet unknown recovery mechanisms [54].

In addition, node strength of the right precuneus was significantly increased in NMDAR encephalitis patients. Previous anatomical and connectivity data suggest a central role for the precuneus in a wide range of higher-order cognitive functions and highly integrated tasks including reflective, self-related processing, emotion-related information processing, and episodic memory [55]. The precuneus is a core region of the DMN and is highly interconnected with the hippocampus [56,57]. It shows reliable increases in activation during both rest and specific tasks and involvement in self-related mental representations during rest. Consequently, it has been proposed that the precuneus is involved in the network correlates of self-consciousness [58]. Hence, structural changes affecting the precuneus may contribute to episodic memory impairment and psychosocial symptoms including decreased judgement of the mental self [55]. However, future studies investigating potential associations between specific neuropsychiatric symptoms such as self-esteem and both functional and structural connectivity of the precuneus are highly warranted.

Our findings complement previous voxel-wise analyses of hippocampal connectivity in 43 NMDAR encephalitis patients that showed reduced functional connectivity between hippocampus and precuneus [6]. Another resting-state fMRI study of seventeen NMDAR encephalitis patients and eighteen matched HC observed decreased amplitude of low-frequency fluctuation (ALFF) in patients in the left precuneus, indicating a decrease in spontaneous neural activity and precuneus functional impairment [59]. In addition, FDG-PET imaging revealed precuneus hypometabolism in six NMDAR encephalitis patients [60]. A recent study using [^18^F]GE-179 PET identified a reduction in the density of open, active NMDARs in the anterior temporal lobes, superior parietal cortices and in the precuneus [61], lending further support to the notion of functional impairment of the precuneus in NMDAR encephalitis.

However, there is only limited data on structural connectivity of the precuneus. Our findings of increased precuneus node strength indicate a relative hyperconnectivity. Indeed, recently discussed mechanisms of compensatory remyelination after inflammatory brain damage might – at least partially – account for increased precuneus node strength in NMDAR encephalitis [62,63].

Alternatively, precuneus node strength increase could be caused by plastic network reorganization given the nature of structural connectome properties and their potential response to acute disease damage. Structural reorganization mechanisms could feature local rerouting that can be thought of as a local outgrowth of new connections due to the diminished capacity of affected hippocampal hubs, previously referred to as ‘hub failure’ [20]. Hence, acute inflammation of central hippocampal hub nodes might lead to a shift from local to global processing to other nodes, i.e. in this case the precuneus. Indeed, previous studies in patients with mild traumatic brain injury support the hypothesis of increased structural connectivity following rerouting of connections from damaged nodes to high-level hubs [64]. Therefore, future investigations in human NMDAR encephalitis are warranted to further elucidate the potential existence of ‘hub failure’ and subsequent hyperconnectivity in NMDAR encephalitis. Moreover, future translational MR studies in murine NMDAR antibody associated disease models [65] may provide additional insights into potential histopathological correlates of these structural hyperconnectivities.

We observed a consistent trend towards lower node strengths in the left and right MTL and a trend for increased mean APL in the left MTL. By contrast, the DMN showed no changes in connectivity on the modular level. Our findings complement previous rs-fMRI analysis that did not detect functional connectivity changes within the DMN itself, but rather a decoupling between DMN and MTL [6]. Correspondence between modular network characteristics in structural and functional connectivity with positive correlations of edge weights between methods have been recently postulated and observed in healthy participants [66,67]. In addition, rs-fMRI based functional connectivity disruption might closely reflect similar structural connectivity degradation in the DMN [68], lending further support to consistent findings across imaging modalities.

As expected, we did not observe differences between NMDAR encephalitis and HC in global measures of network topology. This is in line with findings in schizophrenia that similarly showed nodal and modular structural network changes, including longer node-specific path lengths of bilateral frontal cortex and temporal pole regions, without significant alterations in global network properties [18]. Likewise, functional global network properties were shown to be unaltered in comatose patients when compared with healthy controls [69]. Interestingly, despite the lack of global network alterations in comatose patients, a further in-depth investigation revealed a marked reorganization on the nodal level with reduced hubness of occipital cortex nodes and abnormally increased hubness of nodes in the prefrontal and lateral parietal cortex [69]. Our findings, i.e. the absence of global network alterations in the context of specific alterations on the modular and nodal scale, are in agreement with these previous observations, indicating that modular and nodal changes predominate the structural connectome alterations in in NMDAR encephalitis.

## Limitations

Patients were studied during the post-acute stage of the disease. We, therefore, cannot exclude the presence of global network changes during the acute stages of NMDAR encephalitis. Future longitudinal studies during the early and later stages of the disease are needed to further elucidate the patterns of network alterations in NMDAR encephalitis over time. DWI acquisition was limited to a b-value of 1000, exclusively allowing for single-tissue CSD model creation. While CSD-based probabilistic tractography still outperforms deterministic variants (Farquharson et al., 2013) at this level, a b-value of 3000 with subsequent multi-shell multi-tissue CSD has been suggested to further minimize detrimental effects on tractogram construction (Tournier et al., 2013).

## Conclusion

We employed advanced tractography and graph-theoretical methods to investigate structural connectivity networks in NMDAR encephalitis. Our results reveal that medial temporal lobe structures, specifically the hippocampus, exhibit impaired connectivity, while higher-level network topology remains unaffected. Our study provides further evidence for the specific vulnerability of the hippocampus in NMDAR encephalitis, leading to a critical network hub failure. The correlation of hippocampal node strength with verbal memory performance suggests that structural hippocampal graph metrics may serve as potential MRI markers for assessing cognitive function in the post-acute stage of the disease. Future studies in larger NMDAR encephalitis populations at the acute and post-acute disease stage are warranted to evaluate the clinical utility of diffusion-weighted imaging-based structural connectivity and graph theoretical analysis for disease monitoring in individual patients.

## Acknowledgments

We thank Susan Pikol and Cynthia Kraut for their excellent technical support.

## Funding

JKu was participant in the BIH Charité (Junior) (Digital) Clinician Scientist Program funded by the Charité – Universitätsmedizin Berlin, and the Berlin Institute of Health at Charité (BIH).

## Conflicts of interest

Nothing to report.

## List of abbreviations

ALFF: amplitude of low-frequency fluctuation
APL: average shortest path length
BC: betweenness centrality
CC: clustering coefficient
CMFG: caudal middle frontal gyrus
CNS: central nervous system
CSD: constrained spherical deconvolution
DMN: default mode network
DTI: diffusion tensor imaging
DWI: diffusion-weighted imaging
EC: entorhinal cortex
FA: fractional anisotropy
FLIRT: FMRIB’s Linear Image Registration Tool
FOV: field of view
HC: healthy controls
HI: hippocampus;
MD: mean diffusivity
MS: multiple sclerosis
MTL: medial temporal lobe
NMDAR encephalitis: anti-N-methyl-D-aspartate receptor encephalitis
PC: participation coefficient (PC)
PCU: precuneus;
ROI: region-of-interest
rs-fMRI: resting state functional MRI
TE: echo time
TI: inversion time
TR: repetition time

## Notes

### Competing Interest Statement

The authors have declared no competing interest.

