## Supplementary Material for "Hippocampal hub failure is linked to long-term memory impairment in anti-NMDA-receptor encephalitis - Insights from structural connectome graph theoretical network analysis"

#### *Nodular und Modular Graph Metrics of all nodes for all parameters*

**Supplementary Table S1.** Node Strength for every MTL and DMN region

Values are displayed as mean ± standard deviation. Use of L. and R. denotes left and right. respectively

| Anatomical Region | HC | | | NMDA | | | T-value | p-value |
|---|---|---|---|---|---|---|---|---|
| **L.EC** | $1.67 \cdot 10^8$ | ± | $0.42 \cdot 10^8$ | $1.59 \cdot 10^8$ | ± | $0.36 \cdot 10^8$ | 1.070 | 0.281 |
| **L.HI** | $3.64 \cdot 10^8$ | ± | $0.51 \cdot 10^8$ | $3.45 \cdot 10^8$ | ± | $0.51 \cdot 10^8$ | 2.021 | 0.049* |
| **L.PHIG** | $1.73 \cdot 10^8$ | ± | $0.27 \cdot 10^8$ | $1.70 \cdot 10^8$ | ± | $0.30 \cdot 10^8$ | 0.506 | 0.609 |
| **R.EC** | $1.43 \cdot 10^8$ | ± | $0.34 \cdot 10^8$ | $1.46 \cdot 10^8$ | ± | $0.35 \cdot 10^8$ | -0.409 | 0.682 |
| **R.HI** | $3.44 \cdot 10^8$ | ± | $0.42 \cdot 10^8$ | $3.26 \cdot 10^8$ | ± | $0.39 \cdot 10^8$ | 2.479 | 0.013* |
| **R.PHIG** | $1.69 \cdot 10^8$ | ± | $0.24 \cdot 10^8$ | $1.63 \cdot 10^8$ | ± | $0.26 \cdot 10^8$ | 1.311 | 0.192 |
| **L.CACG** | $1.41 \cdot 10^8$ | ± | $0.28 \cdot 10^8$ | $1.42 \cdot 10^8$ | ± | $0.27 \cdot 10^8$ | -0.160 | 0.870 |
| **L.CMFG** | $2.56 \cdot 10^8$ | ± | $0.36 \cdot 10^8$ | $2.52 \cdot 10^8$ | ± | $0.28 \cdot 10^8$ | 0.595 | 0.540 |
| **L.IPG** | $3.57 \cdot 10^8$ | ± | $0.49 \cdot 10^8$ | $3.64 \cdot 10^8$ | ± | $0.43 \cdot 10^8$ | -0.834 | 0.410 |
| **L.MOFG** | $3.76 \cdot 10^8$ | ± | $0.71 \cdot 10^8$ | $3.83 \cdot 10^8$ | ± | $0.61 \cdot 10^8$ | -0.514 | 0.614 |
| **L.PCG** | $1.69 \cdot 10^8$ | ± | $0.27 \cdot 10^8$ | $1.72 \cdot 10^8$ | ± | $0.25 \cdot 10^8$ | -0.600 | 0.549 |
| **L.RACG** | $1.95 \cdot 10^8$ | ± | $0.33 \cdot 10^8$ | $2.02 \cdot 10^8$ | ± | $0.37 \cdot 10^8$ | -1.078 | 0.282 |
| **L.RMFG** | $4.71 \cdot 10^8$ | ± | $0.73 \cdot 10^8$ | $4.79 \cdot 10^8$ | ± | $0.61 \cdot 10^8$ | -0.618 | 0.537 |
| **L.PCU** | $4.06 \cdot 10^8$ | ± | $0.56 \cdot 10^8$ | $4.20 \cdot 10^8$ | ± | $0.42 \cdot 10^8$ | -1.635 | 0.100 |
| **R.CACG** | $1.49 \cdot 10^8$ | ± | $0.24 \cdot 10^8$ | $1.50 \cdot 10^8$ | ± | $0.22 \cdot 10^8$ | -0.412 | 0.685 |
| **R.CMFG** | $2.34 \cdot 10^8$ | ± | $0.35 \cdot 10^8$ | $2.36 \cdot 10^8$ | ± | $0.22 \cdot 10^8$ | -0.254 | 0.799 |
| **R.IPG** | $4.14 \cdot 10^8$ | ± | $0.49 \cdot 10^8$ | $4.13 \cdot 10^8$ | ± | $0.55 \cdot 10^8$ | 0.129 | 0.899 |
| **R.MOFG** | $3.04 \cdot 10^8$ | ± | $0.48 \cdot 10^8$ | $3.09 \cdot 10^8$ | ± | $0.49 \cdot 10^8$ | -0.505 | 0.609 |
| **R.PCG** | $1.68 \cdot 10^8$ | ± | $0.26 \cdot 10^8$ | $1.70 \cdot 10^8$ | ± | $0.22 \cdot 10^8$ | -0.621 | 0.541 |
| **R.RACG** | $1.50 \cdot 10^8$ | ± | $0.26 \cdot 10^8$ | $1.59 \cdot 10^8$ | ± | $0.25 \cdot 10^8$ | -1.975 | 0.056 |
| **R.RMFG** | $4.74 \cdot 10^8$ | ± | $0.65 \cdot 10^8$ | $4.79 \cdot 10^8$ | ± | $0.69 \cdot 10^8$ | -0.441 | 0.680 |
| **R.PCU** | $4.02 \cdot 10^8$ | ± | $0.58 \cdot 10^8$ | $4.26 \cdot 10^8$ | ± | $0.50 \cdot 10^8$ | -2.434 | 0.013* |

**Supplementary Table S2.** Nodal Betweenness Centrality for every MTL and DMN region

Values are displayed as mean ± standard deviation. Use of L. and R. denotes left and right. respectively

| Anatomical Region | HC | | | NMDA | | | T-value | p-value |
|---|---|---|---|---|---|---|---|---|
| L.EC | $15.2 \cdot 10^3$ | ± | $20.2 \cdot 10^3$ | $9.5 \cdot 10^3$ | ± | $11.4 \cdot 10^3$ | 1.915 | 0.042* |
| L.HI | $108.7 \cdot 10^3$ | ± | $50.7 \cdot 10^3$ | $98.0 \cdot 10^3$ | ± | $60.9 \cdot 10^3$ | 1.048 | 0.289 |
| L.PHIG | $1.4 \cdot 10^3$ | ± | $6.3 \cdot 10^3$ | $4.5 \cdot 10^3$ | ± | $10.7 \cdot 10^3$ | -1.943 | 0.054 |
| R.EC | $6.7 \cdot 10^3$ | ± | $9.0 \cdot 10^3$ | $7.9 \cdot 10^3$ | ± | $10.9 \cdot 10^3$ | -0.655 | 0.512 |
| R.HI | $92.6 \cdot 10^3$ | ± | $44.1 \cdot 10^3$ | $80.3 \cdot 10^3$ | ± | $49.4 \cdot 10^3$ | 1.437 | 0.151 |
| R.PHIG | $5.2 \cdot 10^3$ | ± | $12.5 \cdot 10^3$ | $2.5 \cdot 10^3$ | ± | $5.5 \cdot 10^3$ | 1.543 | 0.126 |
| L.CACG | $2.6 \cdot 10^3$ | ± | $11.8 \cdot 10^3$ | $13.6 \cdot 10^3$ | ± | $49.7 \cdot 10^3$ | -1.675 | 0.108 |
| L.CMFG | $10.5 \cdot 10^3$ | ± | $14.8 \cdot 10^3$ | $5.3 \cdot 10^3$ | ± | $12.2 \cdot 10^3$ | 2.105 | 0.037* |
| L.IPG | $98.3 \cdot 10^3$ | ± | $59.7 \cdot 10^3$ | $102.4 \cdot 10^3$ | ± | $59.5 \cdot 10^3$ | -0.383 | 0.703 |
| L.MOFG | $130.7 \cdot 10^3$ | ± | $126.6 \cdot 10^3$ | $128.3 \cdot 10^3$ | ± | $127.9 \cdot 10^3$ | 0.103 | 0.919 |
| L.PCG | $22.9 \cdot 10^3$ | ± | $23.1 \cdot 10^3$ | $21.0 \cdot 10^3$ | ± | $24.8 \cdot 10^3$ | 0.422 | 0.685 |
| L.RACG | $5.0 \cdot 10^3$ | ± | $13.7 \cdot 10^3$ | $9.6 \cdot 10^3$ | ± | $20.9 \cdot 10^3$ | -1.417 | 0.172 |
| L.RMFG | $163.2 \cdot 10^3$ | ± | $70.2 \cdot 10^3$ | $161.6 \cdot 10^3$ | ± | $71.7 \cdot 10^3$ | 0.128 | 0.895 |
| L.PCU | $120.4 \cdot 10^3$ | ± | $93.8 \cdot 10^3$ | $120.6 \cdot 10^3$ | ± | $110.2 \cdot 10^3$ | -0.011 | 0.992 |
| R.CACG | $3.3 \cdot 10^3$ | ± | $7.6 \cdot 10^3$ | $13.8 \cdot 10^3$ | ± | $49.5 \cdot 10^3$ | -1.630 | 0.130 |
| R.CMFG | $5.9 \cdot 10^3$ | ± | $24.5 \cdot 10^3$ | $10.9 \cdot 10^3$ | ± | $33.4 \cdot 10^3$ | -0.936 | 0.337 |
| R.IPG | $188.1 \cdot 10^3$ | ± | $63.0 \cdot 10^3$ | $169.0 \cdot 10^3$ | ± | $80.9 \cdot 10^3$ | 1.442 | 0.155 |
| R.MOFG | $112.7 \cdot 10^3$ | ± | $129.6 \cdot 10^3$ | $113.5 \cdot 10^3$ | ± | $130.5 \cdot 10^3$ | -0.036 | 0.972 |
| R.PCG | $15.3 \cdot 10^3$ | ± | $22.4 \cdot 10^3$ | $16.0 \cdot 10^3$ | ± | $20.3 \cdot 10^3$ | -0.196 | 0.843 |
| R.RACG | $2.7 \cdot 10^3$ | ± | $12.0 \cdot 10^3$ | $1.1 \cdot 10^3$ | ± | $3.0 \cdot 10^3$ | 1.041 | 0.402 |
| R.RMFG | $144.3 \cdot 10^3$ | ± | $66.3 \cdot 10^3$ | $141.1 \cdot 10^3$ | ± | $68.9 \cdot 10^3$ | 0.263 | 0.788 |
| R.PCU | $119.6 \cdot 10^3$ | ± | $97.7 \cdot 10^3$ | $121.0 \cdot 10^3$ | ± | $113.0 \cdot 10^3$ | -0.077 | 0.933 |

**Supplementary Table S3.** Nodal Average Path Length for every MTL and DMN region

Values are displayed as mean ± standard deviation. Use of L. and R. denotes left and right. respectively

| Anatomical Region | HC | | | NMDA | | | T-value | p-value |
|---|---|---|---|---|---|---|---|---|
| L.EC | $4.1 \cdot 10^{-4}$ | ± | $0.6 \cdot 10^{-4}$ | $4.3 \cdot 10^{-4}$ | ± | $0.7 \cdot 10^{-4}$ | -1.469 | 0.147 |
| L.HI | $3.3 \cdot 10^{-4}$ | ± | $0.3 \cdot 10^{-4}$ | $3.5 \cdot 10^{-4}$ | ± | $0.5 \cdot 10^{-4}$ | -2.391 | 0.017* |
| L.PHIG | $3.8 \cdot 10^{-4}$ | ± | $0.4 \cdot 10^{-4}$ | $3.9 \cdot 10^{-4}$ | ± | $0.4 \cdot 10^{-4}$ | -1.532 | 0.128 |
| R.EC | $4.4 \cdot 10^{-4}$ | ± | $0.7 \cdot 10^{-4}$ | $4.4 \cdot 10^{-4}$ | ± | $1.0 \cdot 10^{-4}$ | 0.106 | 0.924 |
| R.HI | $3.4 \cdot 10^{-4}$ | ± | $0.4 \cdot 10^{-4}$ | $3.6 \cdot 10^{-4}$ | ± | $0.4 \cdot 10^{-4}$ | -1.953 | 0.052 |
| R.PHIG | $3.9 \cdot 10^{-4}$ | ± | $0.4 \cdot 10^{-4}$ | $4.0 \cdot 10^{-4}$ | ± | $0.4 \cdot 10^{-4}$ | -2.314 | 0.026* |
| L.CACG | $3.6 \cdot 10^{-4}$ | ± | $0.4 \cdot 10^{-4}$ | $3.7 \cdot 10^{-4}$ | ± | $0.4 \cdot 10^{-4}$ | -0.728 | 0.473 |
| L.CMFG | $2.9 \cdot 10^{-4}$ | ± | $0.3 \cdot 10^{-4}$ | $3.0 \cdot 10^{-4}$ | ± | $0.3 \cdot 10^{-4}$ | -1.238 | 0.215 |
| L.IPG | $2.7 \cdot 10^{-4}$ | ± | $0.3 \cdot 10^{-4}$ | $2.7 \cdot 10^{-4}$ | ± | $0.2 \cdot 10^{-4}$ | 0.467 | 0.646 |
| L.MOFG | $3.0 \cdot 10^{-4}$ | ± | $0.4 \cdot 10^{-4}$ | $3.1 \cdot 10^{-4}$ | ± | $0.4 \cdot 10^{-4}$ | -0.596 | 0.547 |

| | | | | | | | | |
|---|---|---|---|---|---|---|---|---|
| **L.PCG** | $3.7 \cdot 10^{-4}$ | ± | $0.4 \cdot 10^{-4}$ | $3.8 \cdot 10^{-4}$ | ± | $0.4 \cdot 10^{-4}$ | -0.713 | 0.466 |
| **L.RACG** | $3.6 \cdot 10^{-4}$ | ± | $0.4 \cdot 10^{-4}$ | $3.6 \cdot 10^{-4}$ | ± | $0.4 \cdot 10^{-4}$ | 0.229 | 0.822 |
| **L.RMFG** | $2.6 \cdot 10^{-4}$ | ± | $0.3 \cdot 10^{-4}$ | $2.6 \cdot 10^{-4}$ | ± | $0.3 \cdot 10^{-4}$ | -0.502 | 0.618 |
| **L.PCU** | $2.8 \cdot 10^{-4}$ | ± | $0.3 \cdot 10^{-4}$ | $2.8 \cdot 10^{-4}$ | ± | $0.2 \cdot 10^{-4}$ | 1.392 | 0.160 |
| **R.CACG** | $3.5 \cdot 10^{-4}$ | ± | $0.4 \cdot 10^{-4}$ | $3.6 \cdot 10^{-4}$ | ± | $0.4 \cdot 10^{-4}$ | -1.439 | 0.150 |
| **R.CMFG** | $3.0 \cdot 10^{-4}$ | ± | $0.3 \cdot 10^{-4}$ | $3.0 \cdot 10^{-4}$ | ± | $0.3 \cdot 10^{-4}$ | -0.328 | 0.747 |
| **R.IPG** | $2.5 \cdot 10^{-4}$ | ± | $0.2 \cdot 10^{-4}$ | $2.5 \cdot 10^{-4}$ | ± | $0.2 \cdot 10^{-4}$ | -0.387 | 0.708 |
| **R.MOFG** | $3.1 \cdot 10^{-4}$ | ± | $0.3 \cdot 10^{-4}$ | $3.2 \cdot 10^{-4}$ | ± | $0.3 \cdot 10^{-4}$ | -1.105 | 0.272 |
| **R.PCG** | $3.8 \cdot 10^{-4}$ | ± | $0.4 \cdot 10^{-4}$ | $3.8 \cdot 10^{-4}$ | ± | $0.4 \cdot 10^{-4}$ | 0.186 | 0.861 |
| **R.RACG** | $4.0 \cdot 10^{-4}$ | ± | $0.4 \cdot 10^{-4}$ | $3.9 \cdot 10^{-4}$ | ± | $0.4 \cdot 10^{-4}$ | 0.447 | 0.654 |
| **R.RMFG** | $2.6 \cdot 10^{-4}$ | ± | $0.2 \cdot 10^{-4}$ | $2.6 \cdot 10^{-4}$ | ± | $0.2 \cdot 10^{-4}$ | -0.844 | 0.396 |
| **R.PCU** | $2.8 \cdot 10^{-4}$ | ± | $0.3 \cdot 10^{-4}$ | $2.7 \cdot 10^{-4}$ | ± | $0.2 \cdot 10^{-4}$ | 1.574 | 0.125 |

**Supplementary Table S4.** Nodal CC for every MTL and DMN region

Values are displayed as mean ± standard deviation. Use of L. and R. denotes left and right. respectively

| **Anatomical Region** | **HC** | | | **NMDA** | | | **T-value** | **p-value** |
|---|---|---|---|---|---|---|---|---|
| **L.EC** | 8.4 | ± | 2.0 | 8.6 | ± | 1.6 | -0.549 | 0.577 |
| **L.HI** | 3.1 | ± | 0.5 | 3.0 | ± | 0.5 | 0.628 | 0.534 |
| **L.PHIG** | 7.2 | ± | 1.7 | 7.5 | ± | 1.6 | -1.116 | 0.258 |
| **R.EC** | 9.3 | ± | 1.9 | 9.3 | ± | 1.9 | 0.167 | 0.875 |
| **R.HI** | 3.1 | ± | 0.5 | 3.0 | ± | 0.5 | 0.306 | 0.763 |
| **R.PHIG** | 6.8 | ± | 1.6 | 7.1 | ± | 1.6 | -1.113 | 0.261 |
| **L.CACG** | 3.6 | ± | 0.6 | 3.6 | ± | 0.6 | -0.576 | 0.561 |
| **L.CMFG** | 4.9 | ± | 1.0 | 5.1 | ± | 0.9 | -1.174 | 0.238 |
| **L.IPG** | 4.3 | ± | 0.8 | 4.3 | ± | 0.7 | 0.041 | 0.969 |
| **L.MOFG** | 5.6 | ± | 1.2 | 5.7 | ± | 1.2 | -0.677 | 0.506 |
| **L.PCG** | 3.1 | ± | 0.6 | 3.1 | ± | 0.5 | 0.053 | 0.960 |
| **L.RACG** | 3.9 | ± | 0.7 | 4.1 | ± | 0.7 | -1.666 | 0.095 |
| **L.RMFG** | 4.9 | ± | 1.1 | 5.1 | ± | 0.9 | -1.061 | 0.298 |
| **L.PCU** | 2.6 | ± | 0.5 | 2.7 | ± | 0.4 | -1.295 | 0.196 |
| **R.CACG** | 3.6 | ± | 0.7 | 3.5 | ± | 0.6 | 0.273 | 0.791 |
| **R.CMFG** | 5.1 | ± | 1.2 | 5.2 | ± | 0.9 | -0.241 | 0.815 |
| **R.IPG** | 4.4 | ± | 0.8 | 4.4 | ± | 0.8 | 0.123 | 0.910 |
| **R.MOFG** | 5.9 | ± | 1.2 | 6.1 | ± | 1.2 | -0.906 | 0.360 |
| **R.PCG** | 3.2 | ± | 0.6 | 3.1 | ± | 0.6 | 0.103 | 0.918 |
| **R.RACG** | 4.0 | ± | 0.8 | 4.1 | ± | 0.7 | -1.257 | 0.204 |
| **R.RMFG** | 4.7 | ± | 1.0 | 4.9 | ± | 0.8 | -1.302 | 0.191 |
| **R.PCU** | 2.6 | ± | 0.5 | 2.7 | ± | 0.5 | -1.044 | 0.296 |

**Abbreviations of Anatomical Regions**

| Abbreviation | Anatomical Region |
|---|---|
| **EC** | entorhinal cortex |
| **HI** | hippocampus |
| **PHIG** | parahippocampal gyrus |
| **CACG** | caudal anterior cingulate |
| **CMFG** | caudal middle frontal gyrus |
| **IPG** | inferior parietal gyrus |
| **MOFG** | medial orbitofrontal gyrus |
| **PCG** | postcentral gyrus |
| **RACG** | rostral anterior cingulate |
| **RMFG** | rostral middle frontal gyrus |
| **PCU** | precuneus |
| **MTL** | medial temporal lobe |
| **DMN** | default-mode network |

**Supplementary Table S5.** Modular MTL and DMN graph metrics in NMDAR encephalitis and HC. Mean (sd) values are given across all subjects per group; HC = healthy controls; anti-NMDAR encephalitis = anti-N-methyl-D-aspartate receptor encephalitis; BCC = betweenness centrality; CC = clustering coefficient; APL = average shortest path length.

| | | HC | | NMDAR encephalitis | | *p*-value |
|---|---|---|---|---|---|---|
| | | *mean* | *sd* | *mean* | *sd* | |
| **Node Strength** | Left MTL | $23.5 \cdot 10^4$ | $4.2 \cdot 10^4$ | $22.5 \cdot 10^4$ | $3.0 \cdot 10^4$ | 0.09 |
| | Right MTL | $21.9 \cdot 10^4$ | $2.5 \cdot 10^4$ | $21.1 \cdot 10^4$ | $2.2 \cdot 10^4$ | 0.09 |
| | Left DMN | $29.6 \cdot 10^4$ | $2.7 \cdot 10^4$ | $30.2 \cdot 10^4$ | $2.1 \cdot 10^4$ | 0.23 |
| | Right DMN | $28.7 \cdot 10^4$ | $2.6 \cdot 10^4$ | $29.3 \cdot 10^4$ | $2.2 \cdot 10^4$ | 0.17 |
| **BC** | Left MTL | 41.8 | 18.5 | 37.3 | 21.7 | 0.23 |
| | Right MTL | 34.8 | 17.2 | 30.2 | 16.7 | 0.14 |
| | Left DMN | 69.2 | 23.6 | 70.3 | 25.0 | 0.80 |
| | Right DMN | 74.0 | 24.3 | 73.3 | 30.4 | 0.90 |
| **CC** | Left MTL | $6.2 \cdot 10^{-3}$ | $1.2 \cdot 10^{-3}$ | $6.4 \cdot 10^{-3}$ | $1.1 \cdot 10^{-3}$ | 0.48 |
| | Right MTL | $6.4 \cdot 10^{-3}$ | $1.2 \cdot 10^{-3}$ | $6.5 \cdot 10^{-3}$ | $1.1 \cdot 10^{-3}$ | 0.69 |
| | Left DMN | $4.1 \cdot 10^{-3}$ | $0.7 \cdot 10^{-3}$ | $4.2 \cdot 10^{-3}$ | $0.6 \cdot 10^{-3}$ | 0.34 |
| | Right DMN | $4.2 \cdot 10^{-3}$ | $0.7 \cdot 10^{-3}$ | $4.3 \cdot 10^{-3}$ | $0.6 \cdot 10^{-3}$ | 0.50 |
| **APL** | Left MTL | $3.7 \cdot 10^{-4}$ | $0.4 \cdot 10^{-4}$ | $3.9 \cdot 10^{-4}$ | $0.5 \cdot 10^{-4}$ | 0.05 |
| | Right MTL | $3.9 \cdot 10^{-4}$ | $0.4 \cdot 10^{-4}$ | $4.0 \cdot 10^{-4}$ | $0.5 \cdot 10^{-4}$ | 0.25 |
| | Left DMN | $3.1 \cdot 10^{-4}$ | $0.2 \cdot 10^{-4}$ | $3.2 \cdot 10^{-4}$ | $0.2 \cdot 10^{-4}$ | 0.71 |
| | Right DMN | $3.2 \cdot 10^{-4}$ | $0.2 \cdot 10^{-4}$ | $3.2 \cdot 10^{-4}$ | $0.2 \cdot 10^{-4}$ | 0.75 |

### *Global Graph Metrics*

**Table S6. Network characterization with global and modular graph metrics in NMDAR encephalitis and HC.**

Mean(sd) values are given across all subjects per group with connection densities from 10% to 55%; range values are given as mean(sd) for 10% density and 55% density; HC = healthy controls; anti-NMDAR encephalitis = anti-N-methyl-D-aspartate receptor encephalitis; sigma = small-worldness; Eglob = global efficiency; CCglob = global clustering coefficient calculated as the mean CC across all nodes.

| | | HC | NMDAR encephalitis | p-value |
|---|---|---|---|---|
| **Global** | | | | |
| **Sigma** | Mean(sd) | 1.86 (0.74) | 1.87 (0.75) | 0.45 |
| | Range | 3.58 (0.15) – 1.14 (0.04) | 3.60 (0.18) – 1.13 (0.05) | |
| **Eglob** | Mean(sd) | 1.396 (69) | 1.393 (88) | 0.83 |
| | Range | $1.4 \cdot 10^3$ (70) – $1.4 \cdot 10^3$ (70) | $1.4 \cdot 10^3$ (89) – $1.4 \cdot 10^3$ (89) | |
| **CCglob** | Mean(sd) | 0.012 (0.006) | 0.013 (0.006) | 0.30 |
| | Range | 0.031 (0.005) – 0.004 (0.001) | 0.032 (0.004) – 0.004 (0.001) | |
